# Animal Social Network Structures Across the Fast-Slow Continuum

**DOI:** 10.1101/2025.04.10.648003

**Authors:** Ross S. Walker, Andy White, Xander O’Neill, Nina H. Fefferman, Matthew J. Silk

## Abstract

The fast-slow continuum is a key axis of variation in life history strategies that captures the evolutionary trade-off between allocating in lifespan versus reproduction. It has been suggested that social behaviour, and therefore network structure, may vary across the fast-slow continuum, but formal theory remains scarce. We develop a novel mathematical model to examine how the rate of demographic replacement may influence emergent social network structures in natural populations. We additionally consider variation in social preferences and modes of social inheritance. Our key finding is that more rapid demographic replacement can substantially constrain the structure of dynamic social networks. For species with longer generation times, network structures are primarily determined by social preferences, while for shorter generation times they are primarily determined by mechanisms of social inheritance. By considering how demographic replacement can constrain social network organisation, our work provides important insights into social evolutionary ecology.

## Introduction

Life history strategies are defined by trade-offs between survival, development and behaviour, which are in turn subject to constraints imposed by resource availability, environmental conditions, and allometry (Wang et al. 2021; Stearns 2000; Healy *et al*. 2019). A well-known trade-off that shapes life history strategies is the *fast-slow continuum* (Stott et al. 2024; Healy et al. 2019; Stearns 1983, 2000; Promislow & Harvey 1990; Bielby et al. 2007), which describes the relative allocation in lifespan and reproduction, termed *speed of life* (Silk & Hodgson 2021; Stott et al. 2024; Healy et al. 2019; Zipple et al. 2024). Species with slow speed of life are characterised by long lifespans but low reproductive rates (i.e. long generation times). Such species often have greater levels of maternal care and improved juvenile survival rates, e.g., killer whales (*Orcinus orca*) or African elephants (*Loxodonta africana*) (Zipple et al. 2024; Paniw et al. 2018; Weiss et al. 2023; Hoerner et al. 2023; Allen 2006). Conversely, species with fast speed of life have a high reproductive rate but short lifespan (i.e short generation times), often exhibiting higher metabolic rates than slow-lived species (postulated to sometimes have behavioural consequences under the pace of life syndrome hypothesis; Réale *et al*. 2010; Biro & Stamps 2010; Ricklefs & Wikelski 2002; Polverino et al. 2018, although see Royauté et al. 2018). As the fast-slow continuum is a key axis of variation in life history strategies, it is important to understand how it is associated with other characteristics impacting fitness, such as social behaviour. However, few studies have explicitly tested how social behaviour varies across the fast-slow continuum (Salguero-Gómez 2024).

Individual social relationships collectively define the social network structure of a group or population (Krause *et al*. 2014; Sumpter 2005), and this structure can be shaped over time by demographic processes such as replacement (the natural birth and death of individuals) (Shizuka & Johnson 2020). The impact of demographic replacement can be both direct, through removal and addition of possible social partners, and indirect, inducing behavioural changes that shape social dynamics that then modify network structure (Shizuka & Johnson 2020; Ilany & Akçay 2016; Firth et al. 2017). Birth and death rates can also shape social network structure by altering the selective advantage of different social behaviours. For example, if the initial costs of forming a social relationship are high or the benefits are gained gradually over time, then we might expect such relationships to occur more often in species with longer lifespans (Silk & Hodgson 2021). As a species’ position on the fast-slow continuum is closely linked to generation time, it might be expected to influence social network structure and dynamics (Silk & Hodgson 2021). However, covariation between speed of life and social network structure could be a direct outcome of demography, or related to the evolutionary relationships between social behaviour and life history. Consequently, how and why social network structure varies across the fast-slow continuum remains an open question (Silk & Hodgson 2021).

Despite clear links between demography and social structure, demographic processes are rarely incorporated in social network models (Shizuka & Johnson 2020; Pellis *et al*. 2015), with networks typically either static in size and structure (Ferreri et al. 2014; Gallos & Fefferman 2015), or being dynamic but with constant size (Li et al. 2011; Fefferman & Ng 2007b). This restricts the relevance of such studies over longer timescales, where demographic influence on network structure cannot be neglected (Shizuka & Johnson 2020). Theoretical network modelling approaches incorporating more realistic demographic dynamics could be effective tools to better understand how social structure varies across the fast-slow continuum.

Social network structure and dynamics can also be linked to individual-level social preferences (Croft et al. 2016). Diverse preferential behaviours are observed in nature, including for social relationships based on particular traits, such as larger physical size in greater kudu (*Tragelaphus strepsiceros*) (Owen-Smith 1993) or greater maturity in African elephants (McComb *et al*. 2001), or phenotypic similarity (Massen & Koski 2014; Ebenau *et al*. 2019; Seyfarth *et al*. 2012; McPherson *et al*. 2001; Bizzozzero *et al*. 2019; Croft *et al*. 2009). Therefore, models of social structure and dynamics should incorporate preferential behaviour (Brask et al. 2024). Such an implementation was introduced by Fefferman & Ng 2007b. Here, each individual is assigned a social preference and frequently reevaluates their social bonds with respect to their preference, altering their own network accordingly. Social preferences are implemented in the form of network centrality measures, describing each individuals’ network position (Freeman 1978; Sosa et al. 2021), with different measures capturing a range of natural preferential behaviours (Fefferman & Ng 2007b). This work showed that variation in social preferences drives differences in emergent group-level network structures, with simple individual-level mechanisms driving complexity in the network structure. The model has since been extended to address other ecological (Hock et al. 2010; Hock & Fefferman 2011; Greening & Fefferman 2014) and epidemiological (Hock & Fefferman 2012; Gallos & Fefferman 2015; Fefferman & Ng 2007a; Wilson et al. 2020) questions. However, the ecological and evolutionary conclusions drawn are limited since they do not account for demographic processes, which can be important over longer timescales.

A fundamental consideration when integrating demographic replacement into dynamic social network models is *social inheritance*: if and how social relationships are passed from a parent to its offspring (Shizuka & Johnson 2020; Ilany & Akçay 2016; Ilany *et al*. 2021). One model of social inheritance (Ilany & Akçay 2016) suggested that network stability is achieved through individuals socialising with the social circle of their parents, with this model successfully replicating aspects of the observed social network structure of four mammal species. In contrast, parent and offspring social relationships may be disjoint, such as when species exhibit abandonment behaviour (Davenport et al. 2019). Therefore, different forms or extents of social inheritance may also shape long-term animal social dynamics. Recent studies on inheritance patterns in spotted hyenas (*Crocuta crocuta*) (Ilany et al. 2021) and wild baboons (*Papio spp*.) (Roatti *et al*. 2023) highlighted how particular forms of inheritance can have long-lasting influences upon individual social relationships. The role of social inheritance may be especially significant for fast-lived species, if network dynamics are strongly governed by the more rapid replacement of individuals (Shizuka & Johnson 2020; Ilany & Akçay 2016). It is therefore important to consider social inheritance when assessing how demography may constrain the emergence of social network structures (Ilany & Akçay 2016; Shizuka & Johnson 2020).

Here we extend the dynamic social network model of Fefferman & Ng 2007b to include demographic replacement and social inheritance. We consider popularity and closeness social affiliation preferences (as in Fefferman & Ng 2007b), alongside homophilic social preferences (to broaden the range of behaviours captured; Fu et al. 2012). While our model (or *any* model) cannot capture the full diversity of social organisation across the animal kingdom, its clear assumptions allow conceptualisation of the potential for demographic constraints to influence emergence social network structure and can generate hypotheses to be tested with empirical data. We assess whether demographic constraints differ across the fast-slow continuum, modes of social inheritance, and social preference types. Our key finding is that speed of life can constrain social network structure, determining the extent to which particular individual preferences shape social organisation. Further, for shorter generation times social inheritance becomes more important in determining social network organisation. The resulting conceptual relationship between social network structure and generation time yields insights into how demography may constrain and shape co-evolution of life history and sociality.

## Methods

### Initialisation

We model the social network of a population as a directed graph *G* = (*V, E*). Here, *V* is the set of nodes, each representing an individual, and *E* is the set of directed edges representing (not necessarily mutual) social relationships (or bonds) between individuals (social inclinations that underlie patterns of affiliative behaviour, social support or association; Hinde 1976; Moor *et al*. 2024). The graph is initialised with *N* = 100 nodes, each having five outward edges assigned from a uniform distribution across the population. No duplicated edges are allowed, nor edges which connect a node to itself. Additionally, in order to consider homophilous social dynamics, all nodes are assigned a (non-social) phenotype (‘trait’) value, *T*, taken from a truncated normal distribution, 𝒩 (0.5, 0.1), restricted to the interval [0, 1].

### Demography

The demographic process is modelled as a continuous-time Markov chain. We use a birth-death model including density dependence to ensure population resilience around a carrying capacity, and because density-based effects are important for population-level predictions (Hernández *et al*. 2025). We assume density dependence acts only on birth rates, as this reduces the size of the population fluctuations (Supporting Information Section S1.1). Individuals give birth at rate *b* − *qN*, and die at rate *d*. Here, *b* is the maximum birth rate, *N* is the current total population, and *q* = (*b* − *d*)*/K* is the populations’ susceptibility to crowding, with *K* denoting the population carrying capacity. As such, each individual has a probability of giving birth, *P*_birth_, and a probability of death, *P*_death_, which increases and decreases the population size, respectively (see Equations 1-2).

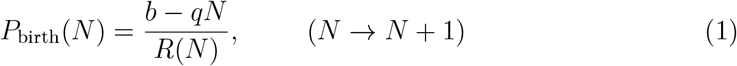

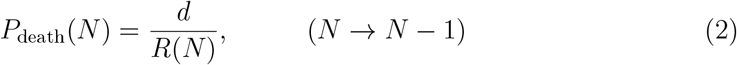

Here, *R*(*N*) = *N* (*b* − *qN*) + *dN* is the sum of the transition rates of the Markov chain at the current state *N*. The time between events is drawn from an exponential distribution with parameter *λ* = *R*(*N*). Note, if *P*_birth_ *<* 0 it is set to zero. This is an application of the Doob-Gillespie algorithm (Doob 1942; Gillespie 1977) for sampling from continuous-time discrete-space Markov chains. Using this process, the average population dynamics are logistic (Caswell 2006), and we assume birth and death processes for each individual are independent.

We capture position on the fast-slow continuum by scaling the birth and death parameters. We define the ‘fastest’ demographic parameters (representing the shortest generation time) as (*b*_*f*_, *d*_*f*_) = (10, 1). Then, by considering a scaling by *c* ∈ [0, 1], we use the fastest parameters to obtain a slower speed of life (*b, d*) = (*cb*_*f*_, *cd*_*f*_). A higher value of *c* therefore corresponds to a faster speed of life. We scale *q* by *c* equally to fix the carrying capacity at *K* = 100. For mathematical details on how *c* correlates with speed of life see Section S1.2, and regarding the choice of values for *b*_*f*_ and *d*_*f*_ see Section S1.3. We note that both generation time and time until first reproduction, established empirical and theoretical measures of position along the fast-slow continuum (Gaillard et al. 2005), vary *inversely* with *c* (Section S1.4).

### Social Inheritance

To integrate birth processes into the network structure, our model includes social inheritance. Upon the birth of a new individual, we consider three different modes of social inheritance: (i) *Zero-inheritance*, where no edges are assigned to the newborn at birth, (ii) *Full-inheritance*, where newborns are assigned the same outward edges as their parent, and (iii) *Parental-inheritance*, where newborns are given an outward edge to their parent only. Whenever an individual dies, their corresponding node is removed from the graph, along with all their edges.

We also include trait inheritance. The new node (individual) is assigned the trait value *T*_*o*_ ∼ *N* (*T*_*p*_, *m*), where *T*_*p*_ is the trait value of the parent and *m* = 0.25 is the mutation variance. Trait values are fixed to the range [0, 1] using reflective boundary conditions. The value *m* = 0.25 was chosen so genetic drift does result in the distribution of trait values becoming narrower during simulation time.

### Social Dynamics

To model social dynamics we follow Fefferman & Ng 2007b. The dynamics are composed of multiple *social updates*, occurring periodically. These are driven by preferential behaviour, assuming that individuals act selfishly to maximise the quality of their own relationships (edges) according to one of three social rules. Prior to the social update step, each node in the graph can have between 0 and 5 outward edges. Nodes with greater than 3 edges may improve the quality of their relationships by dropping their worst scoring ones based on a particular rule (their social preference type), until exactly 3 remain. Then, all nodes form new outward edges to other nodes, chosen uniformly from *V* but excluding their current and recently removed edges, until each individual has 5 out edges. We assume the frequency of social updates is independent of generation time.

Individuals are scored using the individual-level scores given in Table 1, with each corresponding to a social preference. Our three social preferences are: (i) Popularity, where a social relationship is preferred if it has a high in-degree (high number of incoming edges), (ii) Closeness, where a social relationship is preferred if that individual has high closeness centrality (is well integrated socially with a strong ability to reach others within the network), and (iii) Homophily, where a social relationship is preferred if with an individual of a similar trait value. In each simulation, we assume all individuals have the same social preference. For homophily, we additionally consider the scenario where relationships added during a social update are those that are the most preferred from the remaining population (including individuals removed when dropping the lowest scoring relationship), to consider a scenario where social preferences are based on an easily observed and assessed phenotype. We refer to this extra case as ‘complete knowledge’, and the other cases as ‘incomplete knowledge’. These represent two extremes of knowledge in the context of how individuals evaluate their social bonds, which remain biologically realistic (Gaona-Gordillo et al. 2023; Carvalho et al. 2021).

**Table 1.**
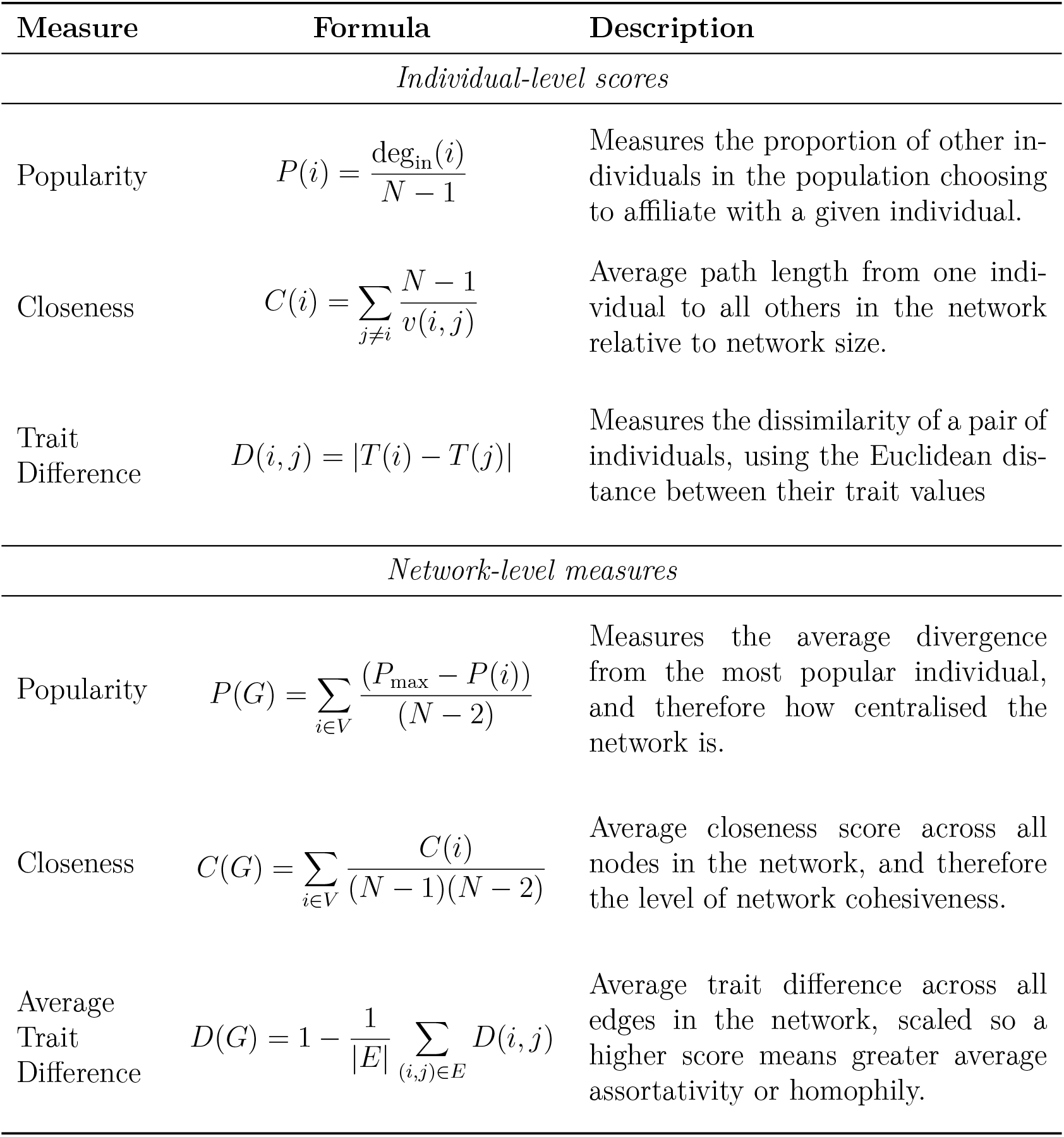
The individual-level and network-level measures considered in this study. Notation: *V* is the set of nodes (individuals), *E* is the set of edges, *T* (*i*) is the trait value of individual *i*, deg_in_(*i*) is the in-degree of individual *i* (meaning the number of edges going *into* this individual), *N* = |*V*| is the total number of individuals, *P*_max_ is the maximum individual-level popularity score in the network and *v*(*i, j*) is the length of the shortest path from individual *i* to individual *j* (where *v*(*i, j*) = *N* if no path exists). The popularity and closeness measures here are as defined in Fefferman & Ng 2007b. Individual measures drive social updates and global measures evaluate organisational success and describe structural properties of the network.

### Model Overview and Simulation

After initialisation, we run the demographic process for *t* = *t*_*D*_. Within this process if a new individual is born, we implement a mode of social inheritance. If a death occurs, all edges for that individual are removed. After this time of *t* = *t*_*D*_ has elapsed, a ‘social update’ is implemented, and the demographic processes continue for another duration of *t* = *t*_*D*_. This process is repeated for a sufficient period of time for the social network structural properties to converge to a quasi-stable state (500 social updates). If *c* = 0 our model is identical to that of Fefferman & Ng 2007b, which is therefore a null model to our own. Each component of the full model is visualised in Figure 1. We choose *t*_*D*_ = 1 without loss of generality (see Section S1.6)

**Figure 1.**
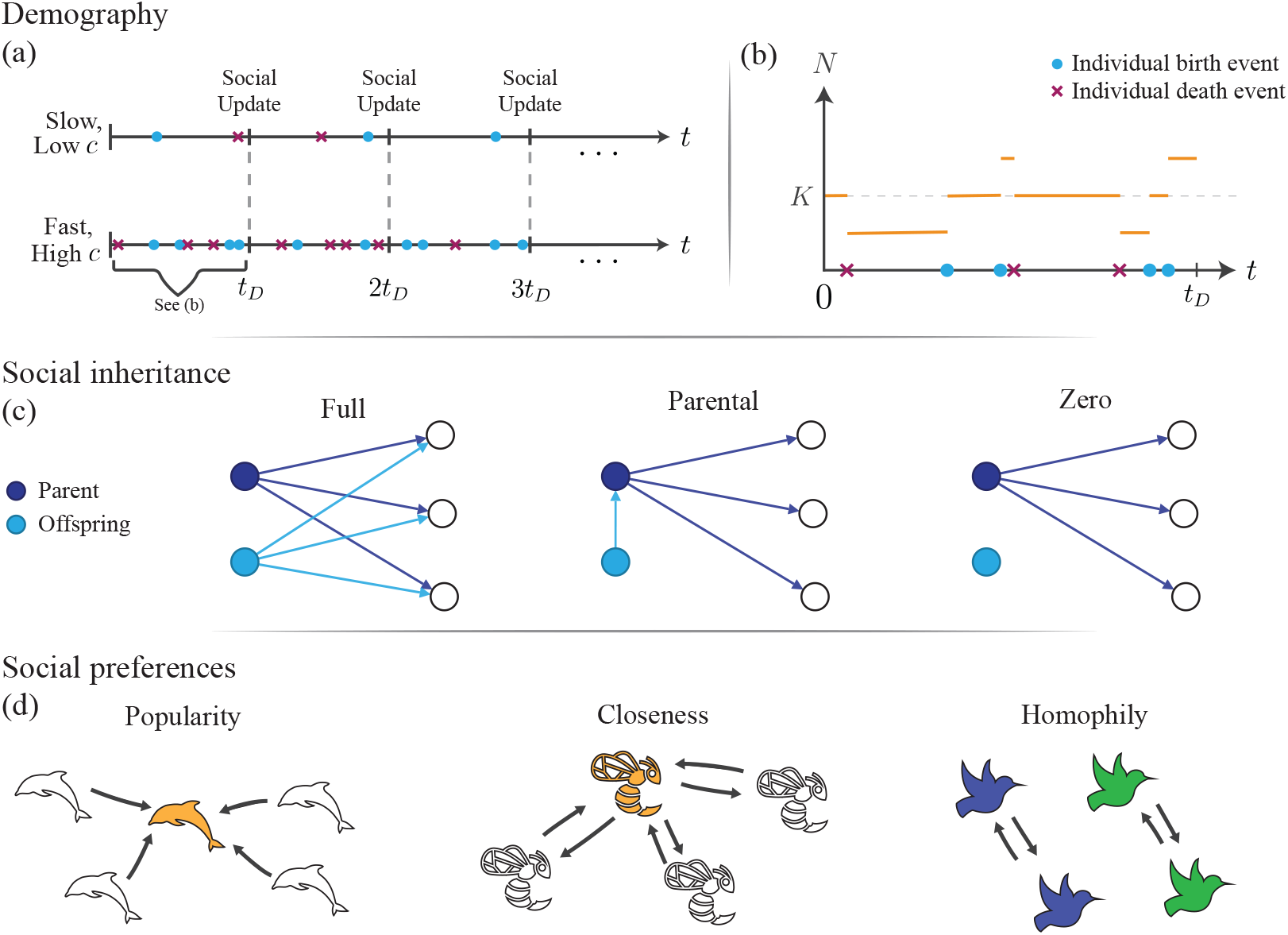
Graphical representation of the model components. (a) Timeline of the model, highlighting the position of the social updates alongside birth (blue dots) and death (purple crosses) events, with (top) high (bottom) low generation times. (b) The changing population size between one social update, as a realisation of a continuous-time Markov Chain model. (c) The different social inheritance schemes, with a parent (dark blue), individuals they are connected to (white) and their offspring (light blue) alongside the relationships assigned at birth. (d) The different social preferences, with high scoring individuals highlighted in orange. For homophily preferences, individuals of similar traits aim to connect with each other (dark blue with dark blue, and green with green).

We quantify differences in social structure with varying generation time by recording network-level measures (Table 1) after each social update. These measures indicate to what extent the social organisation of the group or population is achieved according to one of three social preference rules: popularity, closeness and homophily. Across the fast-slow continuum (100 values of *c*), we perform 100 simulations for each social inheritance mode and each social preference type in order to analyse how the network-level scores change with time, totalling 120000 simulations. The recorded network-level measures are then taken as the mean (over the 100 simulations) of the final 100 social updates. Further details of the simulations are provided in Section S1.5.

## Results

Our simulations provide results relating speed of life (through generation time), the mode of social inheritance, and social preferences through their collective influence upon emergent social network structure. These showed that changes in generation time can substantially impact dynamic social network structure, with this relationship dependent upon both social inheritance and the social preferences used by individuals to update their networks. The results for incomplete knowledge demonstrated that when generation times are long, emergent social network structures are primarily determined by social preferences, whereas when generation time is short social inheritance played a larger role. This is because demographic replacement is typically more rapid in fast-lived species, providing fewer opportunities for individuals to update their networks based on their social preferences. When individuals are equipped with full knowledge, social dynamics were more robust to changes in generation time.

### Popularity Preferences

When social dynamics were driven by popularity preferences, there was substantial variation in network structure across the fast-slow continuum. In our baseline model, with the absence of demographic replacement (*c* = 0), a highly centralised social network structure with three ‘central hubs’ (Figure 2a) was observed, as in Fefferman & Ng 2007b. However, with a higher rate of demographic replacement (i.e. shorter generation time) the networks became less centralised (Figure 2), but more cohesive. This effect is quantified by the rapid decay in network-level popularity scores and increasing (then saturating) network-level closeness scores as the rate of demographic replacement increases (Figures 3a(i), 3a(ii)). This dramatic variation was due to the more frequent deaths of central hubs in the networks, ultimately resulting in networks being stuck in an unstable state that is attempting to converge towards the structure shown in Figure 2a. Higher morality resulted in networks being stuck in an earlier phase of convergence, with lower centralisation on average. The trait assortativity scores of the observed networks varied little with speed of life, except in the absence of demography (*c* = 0), where the score is anomalously dependent upon initial conditions (Figure 3a(iii)).

**Figure 2.**
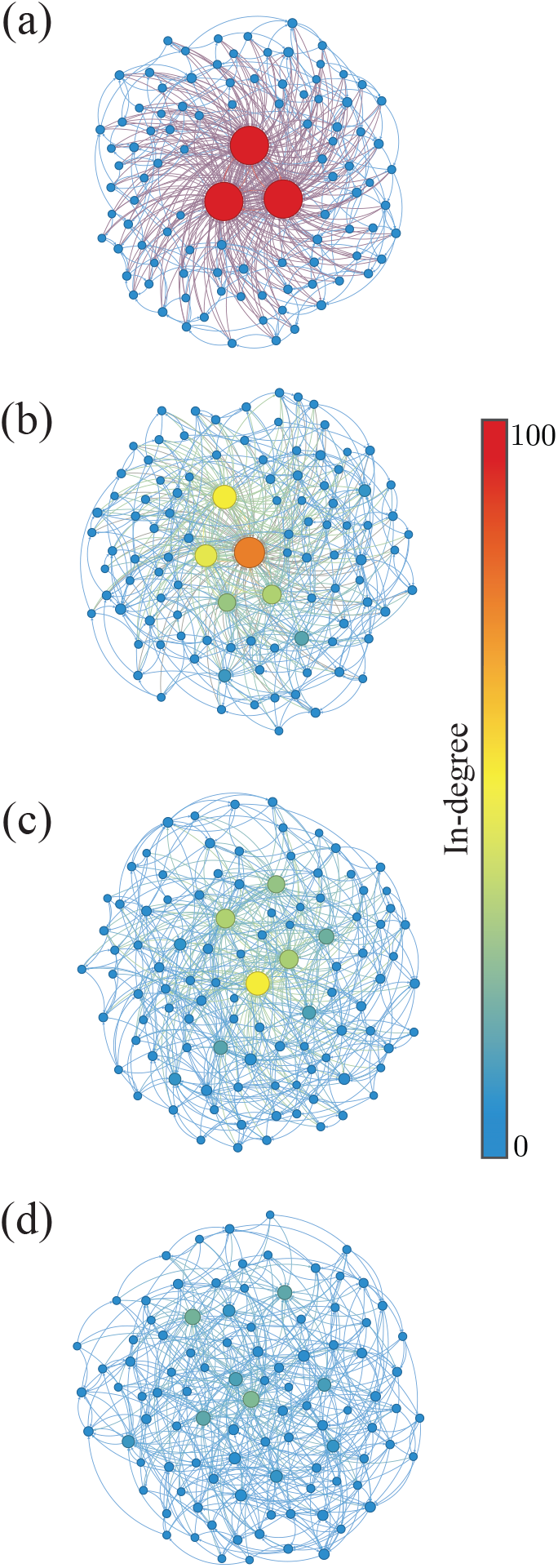
Example social networks obtained as an output of the model for a group with popularity preferences, full social inheritance and replacement rate scaling (a) *c* = 0, (b) *c* = 0.01, (c), *c* = 0.1 and (d) *c* = 0.2. Nodes are sized and coloured according to their in-degree.

**Figure 3.**
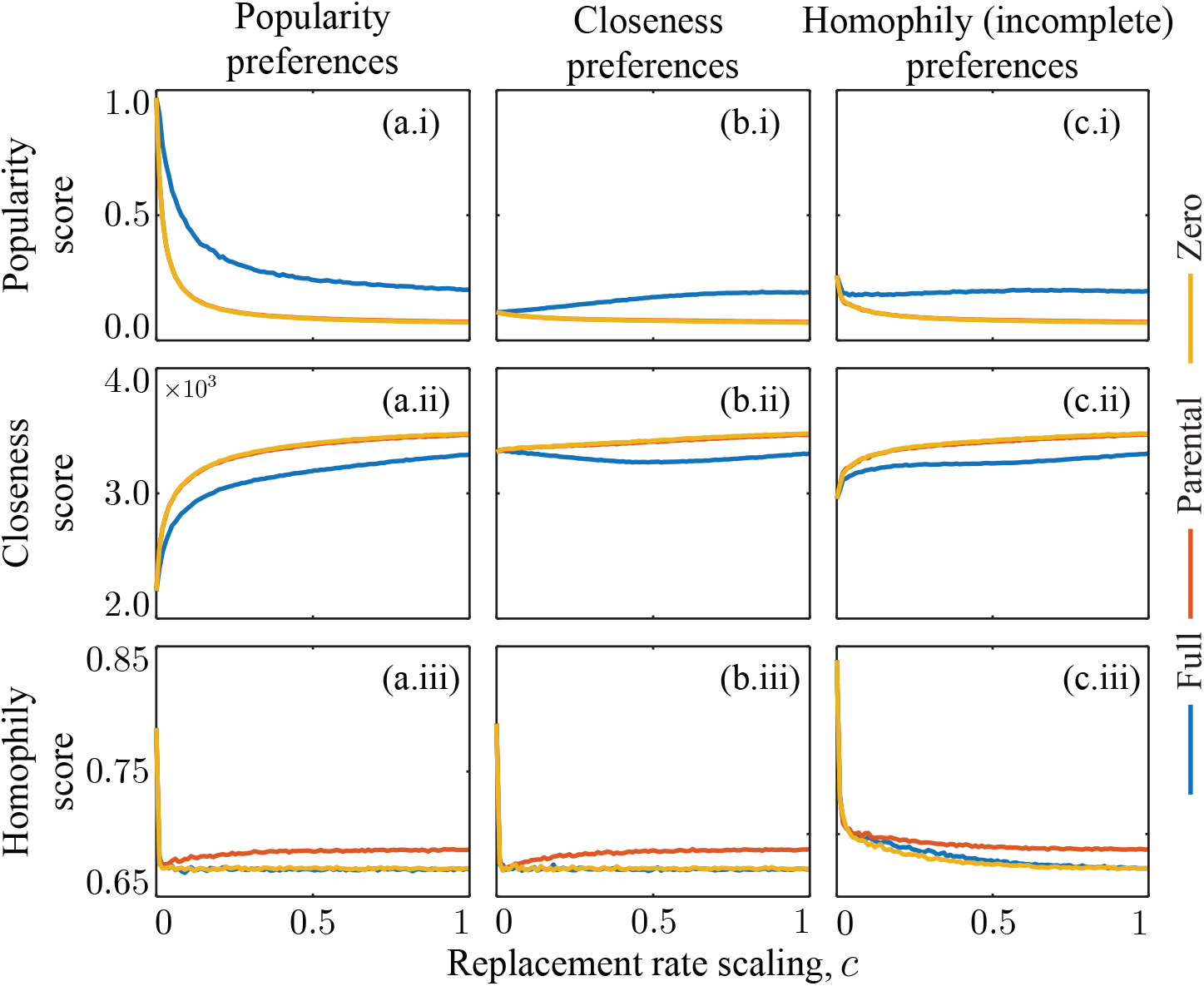
Variation of the average network-level measures with the rate of demographic replacement, represented through variation in *c*. Social preferences are: (a) popularity, (b) *closeness*, and (c) trait (incomplete knowledge). Network-level scores are: (i) popularity, (ii) closeness, and (iii) homophily. Social inheritance modes are: full (blue), parental (red) and zero (yellow). Whenever the parental (red) curve does not appear in a subplot, it had a negligible difference from the zero (yellow) curve.

The mode of social inheritance affected the strength of the relationship between generation time and social network scores by mitigating the impact of the death of central hubs upon network centralisation. With full-inheritance, the magnitude of changes in the structural properties of the network were smaller than with zero or parental-inheritance as full-inheritance led to more rapid replacement of these central hubs. Between zero and parental-inheritance, average network-level popularity and closeness scores were the same. However, parental-inheritance resulted in slightly higher levels of assortativity by trait value, since offspring typically exhibited similar trait values to their parent whom they begin connected to (Figure 3a(iii)).

### Closeness Preferences

When social dynamics were driven by closeness preferences, changes in the rate of demographic replacement caused only small changes in the properties of the network structure (Figure 3b). This invariance might be due to the sensitivity of the closeness measure; rewiring a single edge can dramatically change an individual’s closeness score, making the search for optimal edges difficult. Therefore, the social dynamics themselves are suboptimal under a closeness affiliation preference (Hock *et al*. 2010; Fefferman & Ng 2007b) and less affected by demographic replacement. As with popularity preferences, full-inheritance led to higher centralisation and lower cohesiveness than the other modes of inheritance, and parental-inheritance led to slightly higher trait assortativity. The centralisation of networks slightly increased with the speed of life history, but only with full-inheritance. Here, full-inheritance super-additively makes popular individuals more popular, increasing the group-level popularity score by increasing in-degree heterogeneity. This tendency for full-inheritance to lead to higher popularity scores at faster replacement rates was consistently observed across the other social preferences (Figures 3b-d(i)).

### Trait-based Preferences (Incomplete Knowledge)

When social dynamics were driven by trait-based preferences, and knowledge was incomplete, the effect of variation in demographic replacement rate on the network properties was limited (Figure 3c). As in popularity-driven networks, the change in network-level popularity and closeness scores with increasing demographic replacement was initially rapid, but then saturated. However, the magnitude of the decline was smaller, and there were smaller differences between the network structures of fast-lived and slow-lived species under trait preferences (compared with popularity preferences). The key difference with trait-based preferences is that very high generation times led to a slight increase in network assortativity over the other social preferences (Figure 3c(iii)). Nevertheless, this increase in assortativity score rapidly declined as speed of life increased and generation times were shorter (see Section S2.1).

### Trait-based Preferences (Complete Knowledge)

When individuals had complete knowledge of every individuals’ trait value, the resulting network dynamics and emergent structure were dramatically different to those with incomplete knowledge and, in some circumstances, the emergent social structure was highly organised (Figures 4a-c).

**Figure 4.**
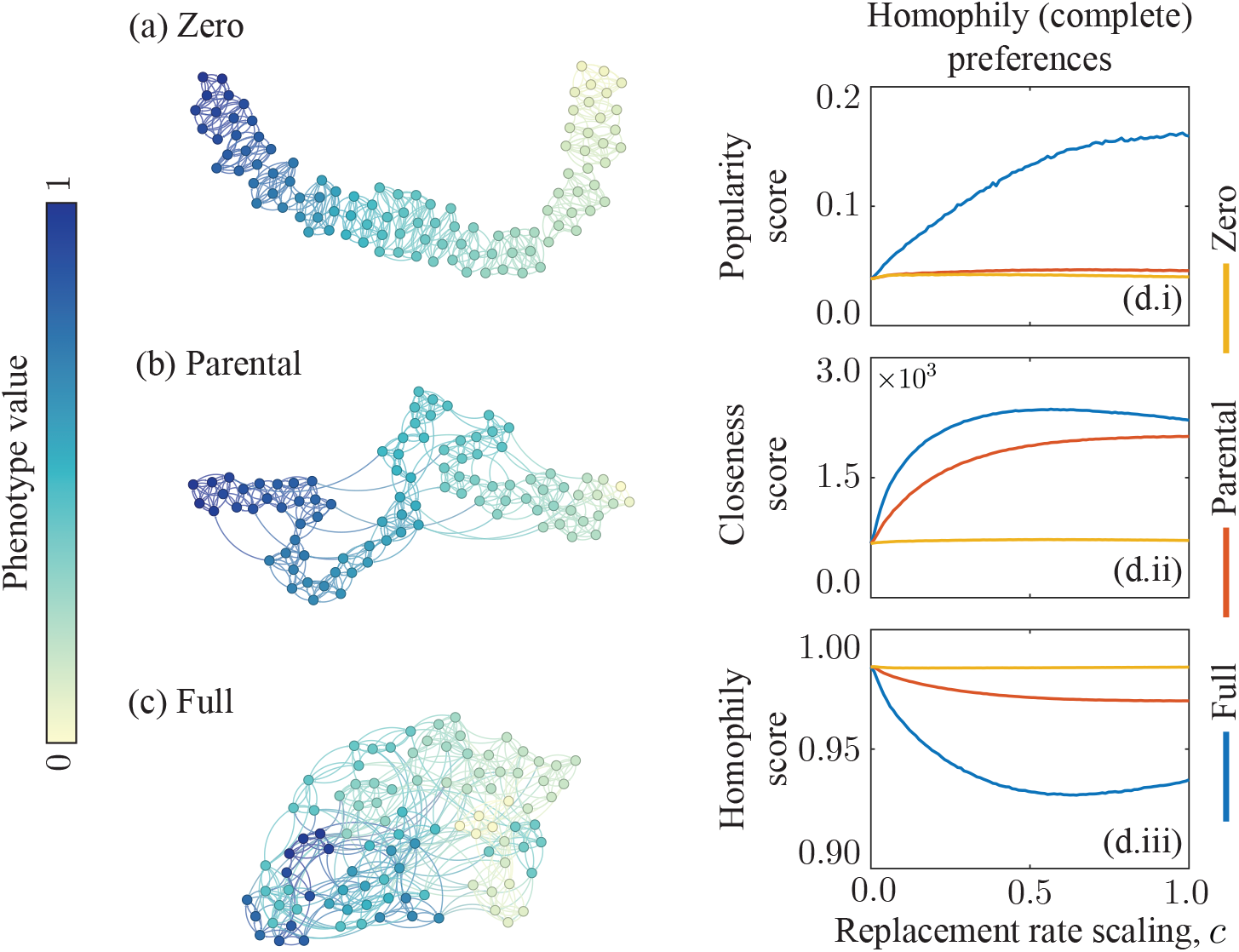
Model results with trait-based (complete knowledge) social preferences. (Left) Example social networks obtained from the model with replacement rate scaling *c* = 0.5 and social inheritance modes: (a) zero, (b) parental, and (c) full. Darker colours correspond to higher trait values. (Right) The variation of the network-level measures with the replacement rate scaling, *c*. Network-level scores are: (i) popularity, (ii) closeness, and (iii) homophily. Social inheritance modes considered are full (blue), parental (red) and zero (yellow).

For trait-based preferences with complete knowledge, popularity and closeness scores were lower than with any social preference with incomplete knowledge (compare Figure 4d(i) & (ii) with Figures 3a-c(i) & 3a-c(ii)). However, complete knowledge led to much higher assortativity scores (compare Figure 4d(iii) to Figures 3a-c(iii)) across the entire fast-slow continuum and all social inheritance modes. In particular, higher social knowledge led to social dynamics being significantly more robust to the impact of demographic processes.

Increasing demographic replacement rate did, however, still impact network dynamics, by increasing the proportion of social bonds formed through inheritance. This is evidenced by network properties being independent to position on the fast-slow continuum with zero social inheritance (Figures 4d(i)-(iii)). With full-knowledge, social dynamics can only add optimal (assortative) edges, while inherited edges may be suboptimal (due to trait mutations). Therefore, without social inheritance there is always an optimal (highly assortative) network structure. Such ‘optimal’ structures (shown in Figure 4a) have low centralisation and cohesiveness. In contrast, with full-inheritance, as rate of replacement increases, more edges are obtained through social inheritance (and these are less strongly assortative). As such, network assortativity decreases while popularity and closeness scores increase (Figure 4c). Parental inheritance balanced network assortativity and cohesiveness; parent-offspring bonds formed bridges across the network between individuals with potentially different trait values, increasing closeness without completely sacrificing assortativity. Homophily is maintained due to the sparsity of these ‘sub-optimal’ social relationships (Figure 4b). Note this effect depends on trait heritability; reducing *m* (increasing heritability) would cause results for parental and full-inheritance to be closer to those for zero-inheritance.

## Discussion

Our model showed that differences in species’ position on the fast-slow continuum, represented through differences in generation time, can constrain the structure of animal social networks, with the strength of the constraint depending upon additional factors such as individual social preferences and modes of social inheritance. Our key finding is that stronger emergent social structure could be a more common feature of animals with slow life history, as slower-lived species may have greater capacity for effective self-organisation (aligning with previous work in primates; Demetrius et al. 2024). This is because the network structures of fast-lived species could not converge to ‘optimality’ (where social preferences are well-fulfilled) due to short generation time resulting in few social updates per average lifespan. Hence, these systems may be more dissipative (far from equilibrium; Green et al. 2008). Furthermore, we show that emergent social network structure is primarily determined by social preferences for slow-lived species, while network structure of fast-lived species is primarily influenced by the mode of social inheritance. When generation times are shorter, there are fewer opportunities for individuals to update their relationships based on social preference, so inherited edges have more prominence. Our results also highlight the importance of social knowledge in shaping how phenotypic assortment shapes social organisation. Consequently, our findings allow us to explicitly link generation time with the social structure of natural systems purely through ecological processes.

### Impact of Social Inheritance Across the Fast-Slow Continuum

We found that social inheritance influences emergent social structure, with some patterns consistent across different social preferences. A higher level of social inheritance (e.g., full-inheritance) produced networks with higher popularity scores and lower closeness scores for all considered social preference types (with incomplete knowledge). This relationship implies that greater social inheritance can consistently result in more centralised and less cohesive social structures, with consequences for other ecological processes. For example, centralised networks may result in possible ‘superspreader’ events, where outbreaks of a simple contagion (such as an infectious disease) may be less likely but more explosive when they occur (Lloyd-Smith *et al*. 2005). Although note that *dynamic* social organisation may restrict the impact of these individuals (Hock & Fefferman 2012). Contrastingly, more cohesive social networks have shorter average path lengths between individuals and may therefore spread a complex contagion (such as novel behaviours) more effectively (Firth 2020), and simple contagions may also spread more effectively (although less explosively).

Social inheritance might also influence the relationship between generation time and the properties of the social network. For example, when social dynamics are driven by closeness preferences, the mode of social inheritance determined if some network-level scores were increasing or decreasing with speed of life. Without investigating multiple social inheritance modes, this sophisticated impact of demographic replacement may not have been observed. Therefore, inheritance (of out-edges) plays a key role in social dynamics, suggesting that network modelling studies should explicitly account for various forms of social inheritance whenever demographic processes are included (Ilany & Akçay 2016).

### Role of Knowledge in Social Network Dynamics

Greater social knowledge (of the group as a whole - ‘complete’ - rather than only an individual’s own social network - ‘incomplete’) was found to reduce the impact of demographic replacement on social organisation. With incomplete knowledge we found that replacement restricted assortativity, such that highly structured networks only emerge at long generation times. Therefore, we expect homophily by less visible traits (like personality or social status) to be more common with slow-lived species (where slight organisational success was observed). With complete knowledge the emergent network structure is highly assortative across the entire fast-slow continuum as the social dynamics are more robust to demographic turnover. Therefore, we predict that we might observe homophily by more easily observed phenotypes (like colour) across more systems as these preferences can be more successfully fulfilled in species across the entire fast-slow continuum, represented by the consistently significantly higher homophily scores when knowledge is complete. While both these kinds of homophily occur in nature (physical homophily: Mourier *et al*. 2012; Kelley & Evans 2018, personality homophily: Bhattacharjee et al. 2024; Massen & Koski 2014; Ebenau et al. 2019; Silk et al. 2003; Seyfarth et al. 2012; McPherson et al. 2001), their relative contributions to social structure for species given life histories and ecological contexts remains an open question, and our results provide some clear theoretical predictions to guide empirical work.

When individuals have complete knowledge, greater levels of social inheritance led to more cohesive network structures, but reduced assortativity. Parental inheritance yielded a high cohesiveness with a smaller reduction in assortativity, representing a particularly effective balance between cohesiveness and network assortativity in homophily-driven groups. The balance is particularly effective for fast-lived species, where generation times are short, suggesting that this mode of social inheritance is particularly beneficial in fast-lived systems which benefit from network cohesiveness. This is interesting in comparison to existing theory that parental bonds are more likely in slow-lived systems, due to the significant fitness advantage of parental care (Klug & Bonsall 2014). Collectively, our results for trait-based preferences highlight the potentially key role of information in animal social network organisation (Gokcekus et al. 2021) and draw attention to how this might vary across the fast-slow continuum. Future work may consider the impact of knowledge across different social preferences and modes of social inheritance, and consider how social knowledge is gained across the lifespan.

### Co-evolution of Social Behaviour and Speed of Life

Our model results suggest that variation in social preferences may substantially change the social network structures for slow-lived species, while the network structures of fast-lived species are more invariant to these changes but are strongly influenced by variation in the modes of social inheritance. This broad result was robust across our different forms of social behaviour considered. We therefore postulate that this central result of our model has broad evolutionary consequences (despite the lack of explicit evolutionary processes in the model). Particularly, since we found that position on the fast-slow continuum and social behaviour jointly influence social network structure, we speculate these features might co-evolve to shape network structure.

Individual social network position has been related to fitness both directly (Brask *et al*. 2021; Snyder-Mackler *et al*. 2020; Farine & Whitehead 2015) and indirectly through correlation with fitness-related traits (Pasquaretta *et al*. 2014; McDonald 2007; Silk et al. 2010; Ellis et al. 2017). Group network structure has also been shown to influence individual fitness (Evans et al. 2021; Ballard & Robel 1974; Robson & Traniello 2002; Clarke & Faulkes 1997; Ebensperger et al. 2016). Since social preferences influenced the network properties most prominently at long generation times, we postulate selection pressures on social preferences may be stronger in slow-lived species. This aligns with the hypothesis that in slower-lived species, (reciprocal) social behaviours may be more evolutionarily viable (and therefore subject to selection on, say, the form of this behaviour), as the costs of these behaviours (including their initial formation) are mitigated by the benefits which accumulate over the longer lifespan (Silk & Hodgson 2021). In contrast, for fast-lived species, selection may be more likely to operate on social behaviours which are more directly related to demography (e.g. inheritance), because demographic processes govern social dynamics more strongly. Conversely, we may also expect varied selection on generation time across different social behaviours. For example, many group-living species benefit from having centralised structures featuring keystone individuals (Ballard & Robel 1974; Knörnschild et al. 2009; Clarke & Faulkes 1997; Robson & Traniello 2002; Brent et al. 2015), which might promote short generation times alongside popularity-driven preferential behaviour (Figure 3a(i)). Yet, these keystone individuals may have different fitness functions (Lloyd-Smith et al. 2005; Aplin et al. 2012).Therefore, social behaviour may co-evolve with generation time, with potential for multi-level selection, due to the collective influence of these features upon the network structure (as uncovered in this study) and the relationship between social structure and fitness (at multiple scales in some contexts). By including explicit evolutionary processes, the co-evolutionary relationships between pace of life history and social behaviour could be directly examined, potentially generating theory shaping our understanding of animal sociality.

## Future Directions

While our model incorporates many key features of natural social system dynamics, it relies upon simplifying assumptions. First, we assume that species update their social relationships at the same rate, independent of lifespan. Assuming a fixed rate of social updating allows *c* to directly correlate with the speed of life history, facilitating straightforward analysis. However, we know little about how social updating varies across the fast-slow continuum (Silk & Hodgson 2021), and pace of life syndromes or co-evolutionary history may break this assumption. Faster-lived species might update social relationships more frequently than slower-lived species due to their greater metabolic rates (Réale et al. 2010) and possibly bolder, more exploratory personalities (Silk & Hodgson 2021). In Section S2.2, we relax this assumption so the rate of socialisation increases or decreases slightly with *c* and show our results are qualitatively robust. It would be interesting to independently vary generation time and age at first reproduction (not currently possible as *c* inversely correlates with both of these features), as this would allow examination of how specific aspects of demography influence emergent social structure.

Additionally, the (uniform) randomness of forming relationships makes it unlikely a simulated network contains multiple connected components. Therefore, our findings are mainly applicable for group-living species (e.g. eusocial insects; Waters & Fewell 2012 or species with a high degree of fission-fusion dynamics; Aureli et al. 2008), and less applicable when we expect greater modularity in social networks. Future work may consider different methodologies for forming relationships, perhaps giving higher weight to second-order social connections in the group, rather than equal weight between all group members. We also assume that our population/group has homogeneous social behaviour - individuals have identical social preferences and aim to keep the same number of relationships. We further assumed, as in Fefferman & Ng 2007b, that the optimal out-degree was 5 and individuals were willing to drop (at most) 2 relationships when adjusting their network. We predict increasing this ratio beyond 5:2 would lead to slower self-organisation (making the emergent structure more sensitive to changes in generation time) but also to stronger emergent structure when the death rate is sufficiently low. It may be interesting to examine if variation in this ratio can lead to *connectivity criticalities* in social network structures (where a large change in the structure arises from a slight variation in behaviour; see Green et al. 2008). Differences or heterogeneities in either the number of social relationships held by individuals or in their social preferences could easily be implemented in future work (Ng 2008; Brask *et al*. 2024). Incorporating individual variation is an interesting direction for future work given the potential importance for ecological processes (Allan et al. 2025; Lloyd-Smith et al. 2005; Stein 2011) and their interaction with social dynamics (Young *et al*. 2022, 2023; Hock & Fefferman 2011).

Here we model varying demography through variation in birth and death rates, neglecting the role of senescence or social ageing. In natural systems the quality and quantity of social relationships can vary across individual lifespans (Sadoughi et al. 2024; Siracusa *et al*. 2022; Wrzus *et al*. 2013). Such effects could be included in our model by implementing weighted edges and a variable ‘optimal’ number of social bonds. If older individuals maintained only their stronger social relationships (Sadoughi et al. 2024), then we may expect increased within-group variation in social position. A weighted implementation of our model could provide exciting new hypotheses on social ageing across the fast-slow continuum. Such implementation could also provide flexibility to model more diverse social behaviours. For example, slow-lived species are hypothesised to have stronger kin-based social bonds (Zipple et al. 2024). This may have evolutionary consequences if extended parent-offspring overlap facilitates broad inter-generational transfers (e.g. of care), which in turn influence life histories. A weighted implementation could capture the effects of differentiated parent-offspring social relationships, providing new insight into the evolution of care.

## Conclusion

Our theoretical model provides key insights into how demographic replacement can constrain social network organisation, and how this might vary with position on the fast-slow continuum. By considering this relationship, our work provides important insights into social ecology and presents hypotheses which may be verified by empirical studies and meta-analyses. Our model is modular so can easily be adapted to consider additional forms of social preferences or dynamics, which may yield predictive potential in specific systems if specific demographic and social parameters are fitted to the model. This work provides new predictions to be tested empirically related to the eco-evolutionary dynamics of social networks, especially as our capability to conduct cross-species comparison of animal social networks grows (Albery *et al*. 2024; Moor et al. 2024). The improved understanding of the link between pace of life history and social network structure that this work provides can help us better understand the drivers of the diverse social life histories observed in nature.

## Acknowledgements

RSW was supported by the EPSRC Centre for Doctoral Training in Mathematical Modelling, Analysis and Computation (MAC-MIGS) funded by the UK Engineering and Physical Sciences Research Council (grant EP/S023291/1). AW and XO were supported by the BBSRC EEID research grant BB/V00378X/1. MJS was supported by Royal Society University Research Fellowship URF/R1/221800. We thank the editor and the three anonymous reviewers for their constructive comments which helped to improve the paper.

