## Supporting Information for "Animal Social Network Structures Across the Fast-Slow Continuum"

#### S. 1 Additional Model Details

##### S. 1.1 Model Density Dependence

We implemented density dependence in our model as a linear decrease in the per-capita rate of reproduction. The deterministic ordinary differential equation (ODE) which describes the average trajectory of these stochastic dynamics is:

$$\frac{dN}{dt} = (b - qN)N - dN. \tag{A}$$

We could also have included density dependence upon the death rate:

$$\frac{dN}{dt} = (b - q_b N)N - (d + q_d N)N. \tag{B}$$

In this section we argue why we chose to exclude density dependence upon the death term. We note that equations (A) and (B) both have the general (logistic) form of

$$\frac{dN}{dt} = N(b - d - xN). \tag{S.1}$$

Fixing the carrying capacity at $N^* = K > 0$ yields the corresponding value of $x$ for each model as

$$x = \begin{cases} q, & \text{Equation (A)} \\ q_b + q_d, & \text{Equation (B)} \end{cases} \tag{S.2}$$

so it is clear that certain parameter choices can make these equations equivalent (besides the trivial case with $q_d = 0$). One key difference between the two models is that, given values of $b, d$, fixing the carrying capacity determines all other parameters (namely, $q$) in Equation (A), while in Equation (B) it imposes the constraint

$$q_b + q_d = \frac{b - d}{K}. \tag{S.3}$$

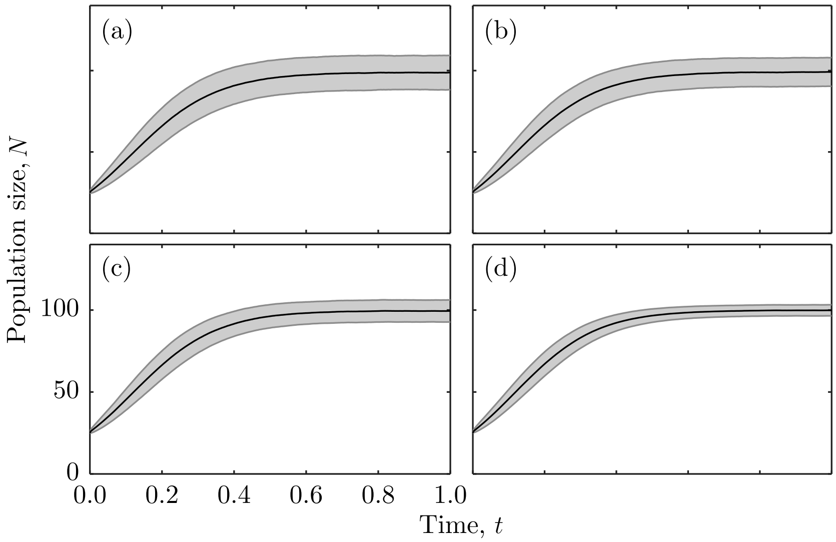


**Figure S.1:** Average population size with time from 10000 stochastic simulations of the demographic model with density dependence upon birth and death. Solid line represents the average trajectory corresponding to Equation (B) and the shaded region represents one standard deviation from the mean trajectory. The parameters are $b = 10$, $d = 1$, $K = 100$, initial population size $N_0 = 25$ and density dependence parameters $q_b, q_d$ determined by Equations (S.4) with (a) $l = 0$ (density dependence only upon death), (b) $l = 0.03$, (c) $l = 0.06$ and (d) $l = 0.09$ (density dependence only upon birth). Average trajectories have decreasing variance with increasing $l$.

Therefore we can introduce the free variable $l \in [0, (b-d)/K]$ such that

$$q_b = l \qquad q_d = \frac{b-d}{K} - l. \tag{S.4}$$

Any value of $l \in [0, (b-d)/K]$ will then fix the carrying capacity as $K$, by construction. Intuitively, $l$ describes the significance of the birth rate density dependence relative to the death rate density dependence. The minimal value of $l = 0$ describes a model with density dependence upon only the death rate, while the maximal model of $l = (b-d)/K$ returns a model with density dependence upon only birth rate (i.e. Equation A). Variation in $l$ does not influence the analytical solutions to (A) due to the $q_b$ and $q_d$ terms summing in the net dynamics (making the ODE independent to $l$). Particularly, the average stochastic realisation of the model will not vary with $l$, and including density dependence upon the death rate will not influence the average trajectory.

However, simulating these dynamics stochastically using the Doob-Gillespie algorithm (as described in the main text), we find that higher values of $l$, corresponding to increased density dependence upon the per-capita death rate, led to increased variance

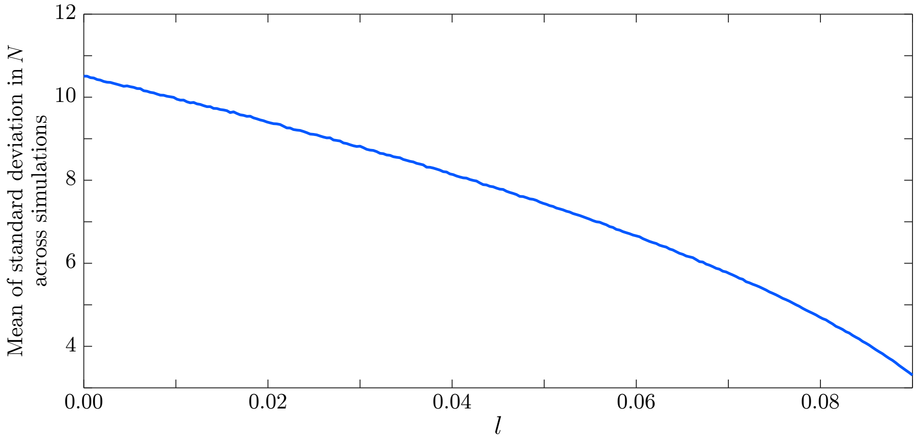


**Figure S.2:** Variation in the standard deviation of the population sizes (averaged across simulation time) with $l$ across 10000 stochastic realisations (per tested $l$ value) of Equation (B). The parameters are $b = 10$, $d = 1$, $K = 100$, initial population size $N_0 = K$ and density dependence parameters $q_b, q_d$ determined by $l$ according to Equations (S.4). Simulations run for period between $t = 0$ and $t = 5$.

in population size across simulations (see Figures S.1 and S.2). Therefore, since our aim is to assess the impact of demographic turnover in a *stable* population upon the network structure, we choose the demographic model with the least variance, namely Equation A, corresponding to $l = (b - d)/K$.

#### S. 1.2 Representing Pace of Life

We capture varying paces of life by scaling the 'fastest' birth and death rates, $b_f, d_f$, by a factor of $c \in [0, 1]$. For $c = 1$ we capture species with fast pace of life, and for $c = 0$ we capture a system with no demographic processes and our model corresponds to that of (Fefferman and Ng 2007). We scale the fastest birth and death rates equally so the ratio of the birth rate and death rate is constant, ensuring that the population remains robust to stochastic extinction across all $c$ values. The state transitions and rates for the Markov chain representing the demographic process are provided in Table S.1.

The time until an event (either a birth or a death), for the parameters representing the fastest pace of life, $T_f$, is taken from an exponential distribution with rate, $R_f(N)$:

$$T_f \sim \exp\left(R_f(N)\right), \tag{S.5}$$

where $R_f(N) = r_{bf}(N) + r_{df}(N)$, and $r_{bf}(N), r_{df}(N)$ are the transition rates for the birth and death events, respectively. The time between events for other (slower) paces

**Table S.1:** Transitions and their respective total rates for the stochastic birth-death simulations, with the fastest and scaled demographic processes considered. Note, the transition rate for birth events is set to zero if $b_f - q_f N < 0$, for avoidance of negative transition rates.

| Parameters | State transition | Transition rates |
|---|---|---|
| $(b_f, d_f)$ | $N \to N+1$ | $r_{b_f}(N) = N(b_f - q_f N)$ |
| | $N \to N-1$ | $r_{d_f}(N) = N d_f$ |
| $(cb_f, cd_f)$ | $N \to N+1$ | $r_{cb_f}(N) = N(cb_f - cq_f N)$ |
| | $N \to N-1$ | $r_{cd_f}(N) = cN d_f$ |

of life, $T_c$, is taken from an exponential distribution with rate, $R_c(N)$:

$$T_c \sim \exp\left(R_c(N)\right), \tag{S.6}$$

where $R_c(N) = r_{bc}(N) + r_{dc}(N)$, and $r_{bc}(N), r_{dc}(N)$ are the transition rates for the birth and death events, respectively. A key observation here is that the expected time between events for species with a slower pace of life, given by $\mathbb{E}(T_c) = 1/R_c(N)$, is proportional to the expected time between events for species with a faster pace of life, given by $\mathbb{E}(t_f) = 1/R_f(N)$:

$$c\mathbb{E}(T_c) = \mathbb{E}(T_f). \tag{S.7}$$

Since the time period for the demographic processes remains fixed ($t_D$ does not vary with $c$), increasing $c$ will increase the expected number of birth and death events in the model, thereby capturing pace of life.

#### S. 1.3 Parameter Choices

We choose $d_f = 1$ so that the shortest average lifespan per individual is one unit of time, i.e. $1/d_f = 1$. We choose $b_f = 10$, the fastest average (maximum) birth rate, to be an order of magnitude greater than $d_f$, making stochastic extinction unlikely and ensuring that the population size does not deviate far from the carrying capacity. We choose a carrying capacity of $K = 100$ so that our demography-free model ($c = 0$) is analogous to the model presented in (Fefferman and Ng 2007). We chose the mutation value, $m = 0.25$, to be large enough that genetic drift does not impact the distribution of trait values. At faster paces of life, smaller values of $m$ resulted in the distribution of trait values becoming extremely narrow. This would maximise the homophily score of the network independent to the social structure, and therefore misrepresent the effectiveness of the social dynamics. The full list of parameters with values and descriptions can be found in Table S.2.

#### S. 1.4 Individual Life History Traits

Our stochastic model of demographic replacement within a population directly defines the expected value of the following life-history traits; generation time, sexual reproductive rate and age at first reproduction.

Since the rate of death events for a given individual is $cd_f$, the time until their death is drawn according to $t \sim \text{Exp}(cd_f)$, so the average lifespan is $1/(cd_f)$. The time between reproductive events (and, due to the memorylessness of the Markov chain model, also the age of first reproduction) is drawn according to $t \sim \text{Exp}(c(b - qN))$, which therefore has expected value $1/(c(b_f - qN))$. Therefore both the average lifespan and the age at first reproduction vary inversely with $c$, but the age at first reproduction also varies inversely with the population size.

When the population is at carrying capacity (so $N = K = \mathbb{E}(N)$), we have that $b_f - qN = d_f$ by definition of $q$. Therefore, at the average population size, the average age at first reproduction is equal to the natural death rate (and therefore the expected number of offspring from a given individual is 1). Particularly, the generation time is the same as the average lifespan, taking value $\frac{1}{cd_f}$. Therefore generation time is also inversely related to $c$.

#### S. 1.5 Analysis of Model Outcomes

For each specific model scenario, we run 100 Monte Carlo simulations. Networks are initialised as outlined in the main paper. The same initial graph structures are used for all variations of the model (all $c$ values, all social preferences, all social inheritance schemes) and each of these initial network structures has a distinct set of initial trait values (randomly drawn from a truncated normal distribution, as outlined in the main

**Table S.2:** Parameter values and their brief descriptions. See main text for full definitions of parameters.

| Parameter | Value(s) | Description | Units |
|---|---|---|---|
| $b_f$ | 10 | Fastest maximum birth rate. | Rate |
| $d_f$ | 1 | Fastest death rate. | Rate |
| $K$ | 100 | Population carrying capacity. | Individuals |
| $m$ | 0.25 | Trait mutation variance. | None |
| $c$ | $[0, 1]$ | Relative speed of demography | None |

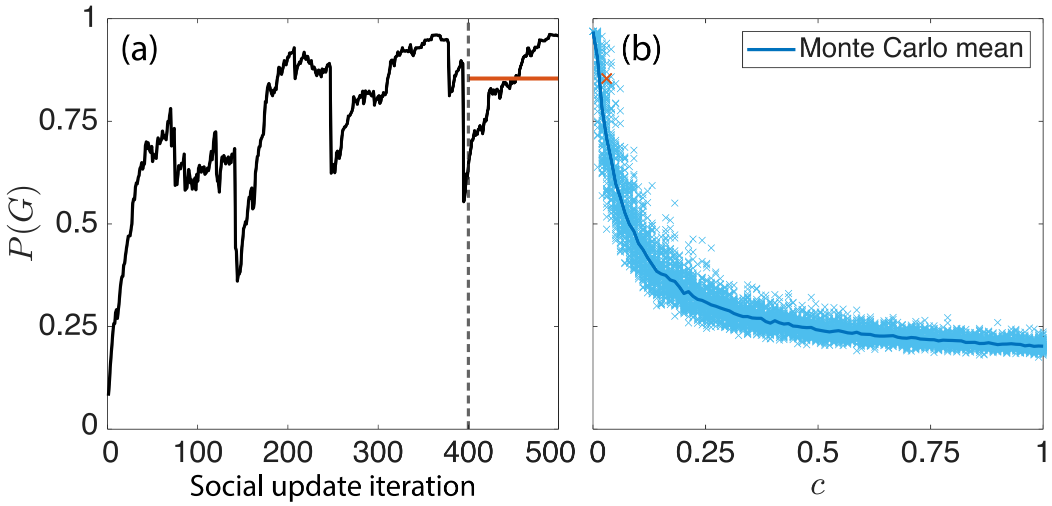


**Figure S.3:** (a) Change in the network-level score for one model realisation with the full-inheritance strategy, popularity social preferences and pace of life $c = 0.0303$ (black), and the mean of the final 100 iterations (red). (b) Final mean values of the 10000 simulations for this particular pair of social inheritance and preference (light blue crosses) alongside the mean curve for each $c$ value (dark blue). The particular simulation from the left is highlighted (red, slightly larger cross).

paper).

In our model simulations, the social networks do not converge towards a fixed final structure, instead the *average* network-level measures begin to saturate with time. Therefore, for each network and simulation, we track the particular measures of interest across the 500 iterations of that particular simulation. We then take the average of the scores across the final 100 iterations as a summary of the final structure (Figure S.3a). The choice of where to average from is purely heuristic. As we perform 100 simulations for each specific scenario, this provides 100 summary scores, which can be averaged to give the final value of each measure for that corresponding $c$ value (Figure S.3b).

#### S. 1.6 Model Generality - Time Between Social Updates

We now show that the choice of the time between social updates of $t_D = 1$ does not affect our findings. This is because changing the value of $t_D$ is mathematically equivalent to changing the pace of life, $c$, and so the results of the model with $t_D = 1$ gives the results of the model for any other value of $t_D$. This ensures that our results regarding the general impact of pace of life on social network properties are independent to this arbitrarily chosen parameter.

To show this result we consider two reparametrisations of the model, where: (1) the time between social updates is rescaled by $\gamma > 0$, but pace of life is fixed, and (2) pace of life is rescaled by $\gamma > 0$, but the time between social updates is fixed. We show that the Markov chains representing the demographic processes in these two scenarios have the same transition probabilities, state space and average number of events between social updates, such that the average changes to the network between social updates are the same for these two scenarios. Throughout, all other model parameters ($b_f$, $d_f$, $K$, social inheritance and social preferences) are fixed.

We denote the (continuous-time, discrete-space) Markov chains representing the demographic processes for the parameter rescalings of $(c, \gamma t_D)$ and $(\gamma c, t_D)$ by $\{N_1(t)\}_{t\geq 0}$ and $\{N_2(t)\}_{t\geq 0}$ respectively. Both Markov chains have an initial distribution of $N_1(0) = N_2(0) = K$ with probability 1 (because simulations are always initialised at the carrying capacity $K$). The key observation is that when both chains are in the same state (meaning $N_1 = N_2 = N$), the transition rates for birth and death events (defined in Table S.3) are proportional:

$$r_{b,1}(N) = \frac{1}{\gamma} r_{b,2}(N), \qquad r_{d,1}(N) = \frac{1}{\gamma} r_{d,2}(N). \tag{S.8}$$

The sum of the transition rates for each of the parameter pairs, denoted $R_1(N)$ and $R_2(N)$ respectively, are therefore also proportional:

$$\begin{aligned} R_1(N) &= r_{b,1}(N) + r_{d,1}(N) && \text{(S.9)} \\ &= \frac{1}{\gamma} \left( r_{b,2}(N) + r_{d,2}(N) \right) && \text{(S.10)} \\ &= \frac{1}{\gamma} R_2(N). && \text{(S.11)} \end{aligned}$$

Therefore, the *relative* transition rates (or the transition probabilities) for the Markov chains for birth ($p_b$) and death ($p_d$) events (from the same state) are equal, since:

$$p_{b,1}(N) = \frac{r_{b,1}(N)}{R_1(N)} = \frac{(1/\gamma) r_{b,2}(N)}{(1/\gamma) R_2(N)} = \frac{r_{b,2}(N)}{R_2(N)} = p_{b,2}(N), \tag{S.12}$$

and

$$p_{d,1}(N) = 1 - p_{b,1}(N) = 1 - p_{b,2}(N) = p_{d,2}(N). \tag{S.13}$$

**Table S.3:** Transition rates in the stochastic birth-death simulation for both $t_D$-scaled and $c$-scaled parameter choices. The transition rate for birth events is set to zero if $b_f - q_f N < 0$, for avoidance of negative transition rates. Note that we are grouping together all birth and death events for all individuals in the network, as to only consider the population-level dynamics.

| **Parameters** | **State transition** | **Transition rates** |
|---|---|---|
| $(c, \gamma t_D)$ | $N_1 \to N_1 + 1$ | $r_{b,1}(N_1) = N_1(cb_f - cq_f N_1)$ |
| | $N_1 \to N_1 - 1$ | $r_{d,1}(N_1) = cd_f N_1$ |
| $(\gamma c, t_D)$ | $N_2 \to N_2 + 1$ | $r_{b,2}(N_2) = N_2(\gamma cb_f - \gamma cq_f N_2)$ |
| | $N_2 \to N_2 - 1$ | $r_{d,2}(N_2) = \gamma cd_f N_2$ |

The transition rates for birth are set to zero whenever the population size is large enough for the birth rate to be negative, giving an upper bound on the population size. Since the birth rates are directly proportional between the two parameter choices (and $\gamma$ is positive), they have the same upper bound on the population size after which further births cannot occur:

$$r_{b,1}(N) \geq 0 \iff r_{b,1}(N) \geq 0 \iff N(cb_f - cq_f N) \geq 0 \iff N \leq \frac{b}{q}. \tag{S.14}$$

Therefore, in both scenarios the maximum population size is given by

$$N_{\max} = \left\lceil \frac{b}{q} \right\rceil. \tag{S.15}$$

Furthermore, the minimum population size is 0, corresponding to extinction (this is the absorbing state of the Markov chain, as the transition rates are all zero at this point). The state spaces of both Markov chains are therefore the same, denoted by $S$ and given in Equation (S.16).

$$S = \{N : N = 0, 1, \ldots, \lceil b_f/q_f \rceil\} \tag{S.16}$$

This implies that the corresponding transition matrices are of the same size, and therefore identical since the transition rates from any given state are also the same.

We now show that the average number of events between social updates is the same for the two Markov chains. The transition times for each Markov chain, denoted by $T_1$ and $T_2$, are drawn from exponential distributions with parameters $R_1(N)$ and $R_2(N)$ respectively:

$$T_1 \sim \text{Exp}(R_1(N)) \qquad T_2 \sim \text{Exp}(R_2(N)) \tag{S.17}$$

and, therefore, the *expected* time between events for the two Markov chains are related according to:

$$\mathbb{E}(T_1) = \frac{1}{R_1} = \frac{\gamma}{R_2} = \gamma \mathbb{E}(T_2). \tag{S.18}$$

Hence, the average number of events between social updates for the chains, denoted by $M_1$ and $M_2$ respectively, are equal:

$$M_1 = \frac{\gamma t_D}{\mathbb{E}(T_1)} = \frac{\gamma t_D}{\gamma \mathbb{E}(T_2)} = \frac{t_D}{\mathbb{E}(T_2)} = M_2. \tag{S.19}$$

Particularly, on average and in a fixed period of time, there are $\gamma$ times as many events with the parameters $(\gamma c, t_D)$ than with the parameters $(c, \gamma t_D)$, although there is $\gamma^{-1}$ times as much time between social updates with parameters $(\gamma c, t_D)$, meaning on average the number of events is the same.

Therefore, as we claimed, the expected population dynamics between social updates with parameters $(c, \gamma t_D)$ are the same as those with $(\gamma c, t_D)$. For example, this tells us that the average model output for $t_D = 2$ and $c = 0.5$ is the same as that of $t_D = 1$ and $c = 1$ (see Figure S.4 for example dynamics between social updates in these cases). Therefore, by considering the range of $c$ values which capture all of the variation in our network measures for a *single* value of $t_D$, these dynamics have also been captured for *any other* choice of $t_D$. We then choose $t_D = 1$ because this yields the neat property that $1/c$ is the number of social updates per average lifespan.

### S. 2 Additional Results

#### S. 2.1 Incomplete Trait Preference Benefit

We can quantify the effectiveness of the social dynamics in the case of incomplete-knowledge trait-based preferences by comparing the average trait similarity score to those observed in networks driven by popularity or closeness preferences. Since these other two network types do not explicitly try to organise by trait values, they give an expected 'random' level of trait assortativity.

We denote by $D_T(c)$, $D_P(c)$ and $D_C(c)$ the average trait similarity (homophily) scores of the network for pace of life $c$ (defined in Table 1 of the main text) when the social dynamics are driven by (incomplete) trait preferences, popularity preferences and closeness preferences respectively. We highlight that the improvement to the assortativity score with the incomplete trait preference, quantified by both of the differences in the homophily scores (Equations (S.20) and (S.21)), decay rapidly with pace of life (Figure S.5).

$$\Delta D_P = D_T(c) - D_P(c) \tag{S.20}$$

$$\Delta D_C = D_T(c) - D_C(c) \tag{S.21}$$

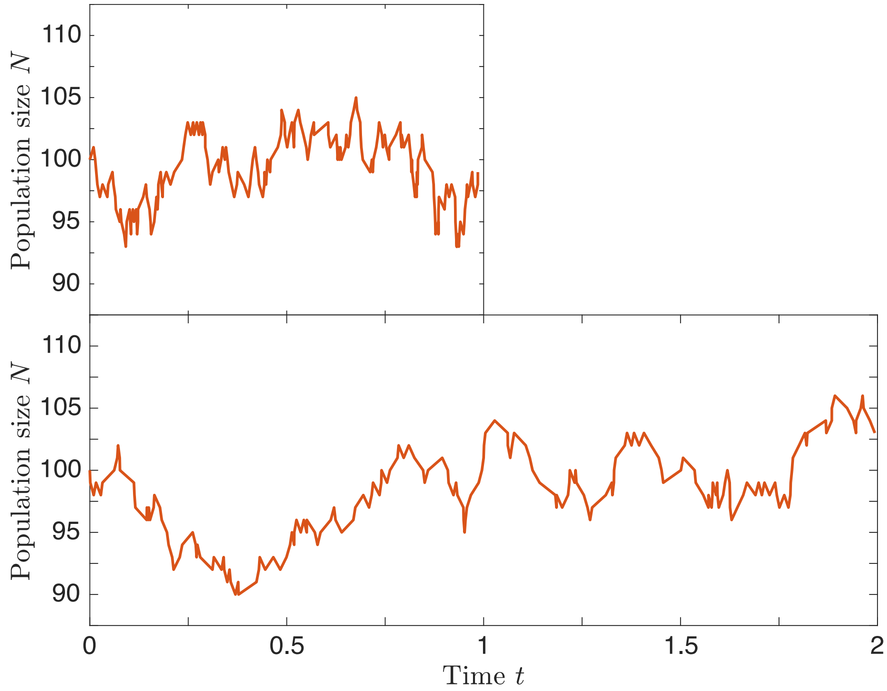


**Figure S.4:** Examples of demographic processes, given as stochastic realisations of the stochastic birth-death model (initialised at the carrying capacity) using the Doob-Gillespie algorithm (Gillespie 1977; Gillespie 2007; Doob 1942; Doob 1945). Fixed parameters are $b_f = 10$, $d_f = 1$ and $K = 100$. Different parameters are (top) $c = 1, t_d = 1$ and (bottom) $c = 0.5, t_d = 2$. The top figure shows more rapid population fluctuations than the bottom figure, but in a shorter time period.

#### S. 2.2 Considering Pace of Life Syndrome

The property uncovered in Subsection S. 1.6 gives the model flexibility, and allows us to easily adjust our results to represent situations where the rate of sociality changes with pace of life. For example, suppose that $t_D = f(c)$ where $f$ is some positive function. This means considering the model outcomes with parameters $(c, t_D) = (c, f(c))$ for $c \in [0, 1]$. We have shown that this is equivalent to considering the model outcomes with parameters $(1, cf(c))$ and then varying $c$.

Let $S(c)$ be the value of one of the network scores (group-level popularity, etc.) obtained from Monte Carlo simulation with parameters $(c, 1)$. Then the value of this same score obtained from Monte Carlo simulation with parameters $(c, f(c))$ should be (neglecting Monte Carlo error) equal to $S(cf(c))$. If $cf(c) > 1$, then we can define $S(cf(c)) = S(1)$, since our initial simulations with $t_D = 1$ covered all $c$ values where there was non-negligible variation (so $S(c)$ for $c > 1$ should deviate little from $S(1)$). This implies that our results with $t_D = 1$ generalise nicely to the cases where $t_D$ is non-constant.

When the rate of sociality decreases (linearly) with pace of life, we find that our current

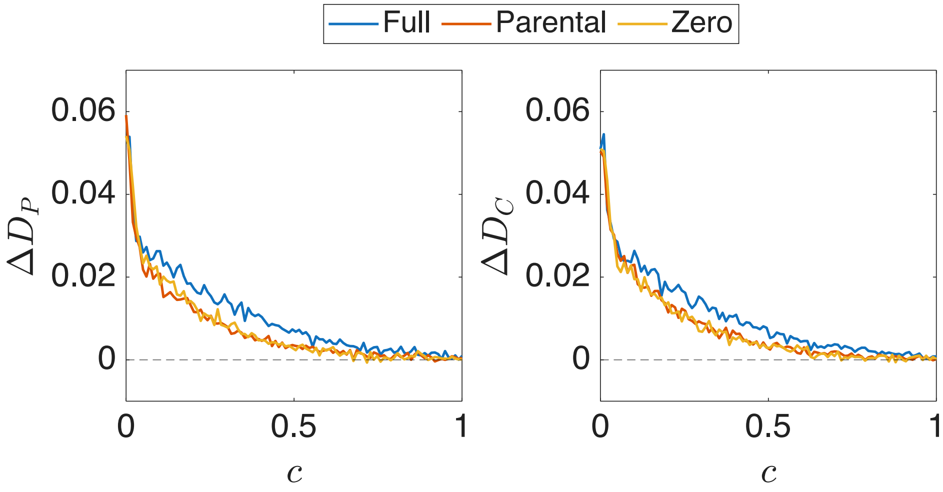


**Figure S.5:** The impact of trait-based social preferences with incomplete social knowledge upon the level of trait homophily of the network, quantified by the values of (a) $\Delta D_p$ and (b) $\Delta D_C$ for $c \in [0, 1]$. Social inheritance strategies considered are *full* (blue), *parental* (red) and *zero* (yellow).

results are qualitatively robust. This is also true when the rate of social updates is increasing slightly with pace of life (possibly representing a weak pace of life syndrome (Silk and Hodgson 2021)). When this relationship becomes more extreme, our results start to change for fast pace of life since the rate of social updates becomes extremely large, and the network structures of extremely slow species are recovered. This is exemplified, for a particular social preference and social inheritance strategy, in Figure S.6 .

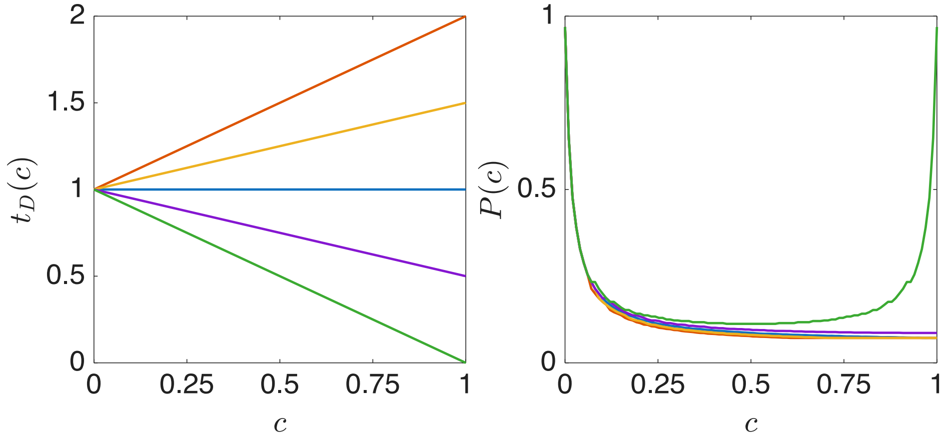


**Figure S.6:** Popularity scores with varying pace of life, $c$, and for different relations between pace of life and rate of sociality, using the relationship outlined in Section S. 2.2. (Left) Different functional relations between pace of life and rate of sociality. (Right) The corresponding change in network-level popularity scores with pace of life. Parameters used are as detailed in Table S.2. The social inheritance strategy is zero-inheritance.
